# Reference genome choice impacts SNP recovery but not evolutionary inference in young species

**DOI:** 10.64898/2026.02.04.703758

**Authors:** Luana S. Soares, Leonardo T. Gonçalves, Sebastián Guzmán-Rodriguez, Aureliano Bombarely, Loreta B. Freitas

**Affiliations:** Department of Genetics, Universidade Federal do Rio Grande do Sul, Porto Alegre, Brazil; University of Nebraska–Lincoln, Lincoln, United States of America; Instituto de Biologia Molecular y Celular de Plantas (IBMCP) (CSIC-UPV), Valencia, Spain

**Author notes:** **Corresponding Author:** E-mail address (L. B. Freitas).

**Keywords:** RAD-seq, rapid divergence, recent speciation, genome choice, bioinformatic approach, population genomics, reference genome

## Abstract

Reduced-representation sequencing approaches such as RAD-seq are widely used in population genomics and phylogenetics, particularly for non-model organisms. However, bioinformatics choices during data processing can strongly influence downstream analyses. One key but underexplored factor is the reference genome used for read alignment and SNP discovery. Here, we evaluate the effects of reference genome choice on RAD-seq analyses using multiple datasets spanning recent radiations in *Petunia* and *Calibrachoa*, and reference genomes that differ in phylogenetic relatedness. When using congeneric reference genomes, we observed highly consistent mapping rates, SNP recovery, and downstream population genomic patterns. In contrast, mapping to more distantly related genomes resulted in lower mapping rates and stronger effects on summary statistics. Despite these quantitative reductions, broader patterns of genetic structure and diversity, as well as evolutionary relationships, remained largely congruent across reference genomes. Overall, our results indicate that reference genome choice matters most when genomes are distantly related or when analyses target fine-scale genomic signals. For recent radiations with largely conserved genome structure, closely related reference genomes yield comparable SNP datasets and lead to the same biological conclusions regarding population structure and phylogenetic relationships. These findings provide practical guidance for RAD-seq studies in non-model systems, showing that congeneric reference genomes are sufficient for robust population and phylogenetic inference, and that more distantly related genomes can remain informative when no close reference is available.

## Introduction

High-throughput sequencing technologies have transformed the study of non-model organisms in ecology and evolutionary biology (Ellegren, 2014; Shafer et al., 2016; 2017). One of the most significant developments has been the advent of reduced-representation sequencing methods, such as restriction-site associated DNA sequencing (RAD-seq). This approach enables the cost-effective and efficient identification of thousands of polymorphic loci across the genome without requiring a complete reference genome (Andrews et al., 2016). RAD-seq has proven particularly useful for population genetics, phylogenetics, and conservation genomics, making it an invaluable tool for researchers working with diverse taxa, including species with poorly annotated genomes.

Despite the advantages of RAD-seq, bioinformatic processing of reduced-representation data presents several challenges and potential sources of bias. These include missing data, which can influence phylogenetic and population genetic inference (Wiens, 1998, 2003; Pybus & Harvey, 2000; Wiens & Moen, 2008; Wiens & Tiu, 2012; Grievink et al., 2013; Lemmon & Lemmon 2013), as well as the effects of bioinformatic filtering on downstream analyses (Slatkin, 1985; Novembre & Stephens, 2008; Nelson et al., 2012; Bhatia et al., 2013; Biddanda et al., 2020; Momozawa & Mizukami, 2021; Weiner et al., 2023). In addition, several studies have shown that filtering out sites with low minor allele frequencies truncates the allele frequency spectrum by removing recent mutations (Shafer et al., 2017; Linck and Battey, 2019), which can bias inferences of historical demography and other evolutionary processes, particularly in the recent past (Boitard et al., 2016).

Beyond filtering and missing data, the strategy used to assemble and genotype RAD-seq data plays a critical role in shaping downstream results. The choice between reference-based and *de novo* assembly approaches can strongly influence SNP discovery, population genetic inference, and evolutionary interpretation (Shafer et al., 2017). Each strategy introduces distinct biases: reference-based approaches can suffer from mapping bias, leading to distorted estimates of heterozygosity and allele frequencies (Günther & Nettelblad, 2019), whereas *de novo* assemblies may result in fragmented loci and elevated sequencing or assembly errors (Davey & Blaxter, 2010). When a high-quality reference genome is available, it can improve SNP recovery, genotype accuracy, inference of demographic history, and the identification of loci under selection (Manel et al., 2016; Brandies et al., 2019). However, for many non-model species, appropriate conspecific references are often unavailable, requiring researchers to either use a reference genome from a closely related species or adopt a *de novo* assembly approach (Reid et al., 2021), a widely recommended approach under the principle of using the “closest available reference genome” (Therkildsen & Palumbi, 2017; Galla et al., 2018). Simulation studies show that even small divergences between the heterospecific reference and the target genome (0.15% – 2%) can bias polymorphism detection and estimates of genetic diversity, particularly at low sequencing depth (Nevado et al., 2014). While empirical studies suggest that closely related genomes may still be suitable for SNP discovery in groups with conserved genomes (Galla, 2019), the degree of evolutionary relatedness required for reliable inference remains ambiguous and likely varies across taxa with different evolutionary histories. This uncertainty is especially pronounced in recently and rapidly diverged species, where shallow divergence and heterogeneous genomic landscapes may amplify reference-based biases. As a result, the impact of reference genome choice in such systems remains unresolved, highlighting the need for further empirical evaluation.

In addition, it has been reported that sequencing depth can influence SNP detection and genotype accuracy (Lin et al., 2024; Liu et al., 2022), potentially compounding the effects of reference choice. Specifically, these reports indicate that, as expected, increasing depth recovers more SNPs up to approximately 20x, after which the discovery rate decreases sharply, signaling saturation of sequencing effort. They point to 10x as the most cost-effective sequencing depth for SNP calling from whole-genome resequencing experiments, and even as low as 6x for population genomics studies, although no study has specifically assessed these constraints for reduced-representation sequencing methodologies such as DArT.

The genera *Petunia* and *Calibrachoa* (Solanaceae) provide an excellent system for investigating this question. *Petunia,* better known for its cultivated *P.* x *hybrida* (Hook.) Vilm., includes 20 wild species that have recently and rapidly diverged (e.g., Särkinen et al., 2013; Reck-Kortmann et al., 2014; Backes et al., 2024; Soares et al., 2025b). Similarly, *Calibrachoa*, a close relative of *Petunia* (Särkinen et al., 2013; Pezzi et al., 2024), comprises species that also exhibit rapid evolutionary divergence (Fregonezi et al., 2012; Mäder & Freitas, 2019; Backes et al., 2025). Although some *Petunia* reference genomes are now available, none exists for *Calibrachoa*. As a result, population genomic studies in *Calibrachoa* must rely on *Petunia* reference genomes (Backes et al., 2025) or employ *de novo* assembly approaches.

In this study, we evaluated how the choice of reference genome affects SNP calling and downstream evolutionary interpretations in *Petunia* and *Calibrachoa*. Specifically, we compared the performance of multiple reference genomes that vary in genetic proximity to the target species to assess how reference selection affects key population and phylogenetic analyses. We hypothesized that greater phylogenetic distance between the reference and focal genomes would reduce mapping quality and SNP-calling accuracy, thereby introducing biases in population genetic inference. Additionally, we tested whether low coverage biases the inference of population genetic statistics. We predicted that low-read-coverage sequencing would have a limited effect on core statistics such as nucleotide diversity (π), inbreeding coefficient (*F*_IS_), or the proportion of polymorphic sites.

By addressing these questions, we provided insights into best practices for conducting genomic analyses of recently and rapidly diverging species. We improved the broader understanding of reference genome selection in population genomics. Instead of focusing on the absolute difference in numbers between the assemblies, we emphasized whether the evolutionary interpretation of the data would differ across tests.

## Materials and Methods

### Study Design and Dataset Overview

To evaluate the impact of reference genome selection on SNP calling and downstream population and phylogenetic inference, we used raw sequencing data from previously published studies, comprising 121 individuals of *Petunia* species (Soares & Freitas, 2024; Soares et al., 2024, 2025a) and 40 individuals of the closely related genus *Calibrachoa* (Backes et al., 2025) (Supplementary Table S1). We structured analyses around three datasets that capture different biological and phylogenetic contexts in which reference genome choice may influence downstream analyses: The POP dataset included 80 individuals from three populations of *P. altiplana*, representing a classical population genetics sampling for a single species. The INTRA-P4 dataset comprised 41 individuals of four *Petunia* species that diverged during the last 2 Mya (Soares et al., 2025a; Figure 1). This dataset enabled assessment of reference-genome effects among closely related congeners that have undergone recent divergence and potential hybridization (Soares et al., 2025a). The INTER-C5 dataset consisted of 40 individuals of five *Calibrachoa* species, a genus that diverged from *Petunia* roughly 7.8 Mya (Huang et al., 2023), providing a test case for reference genome biases when no assembly is available.

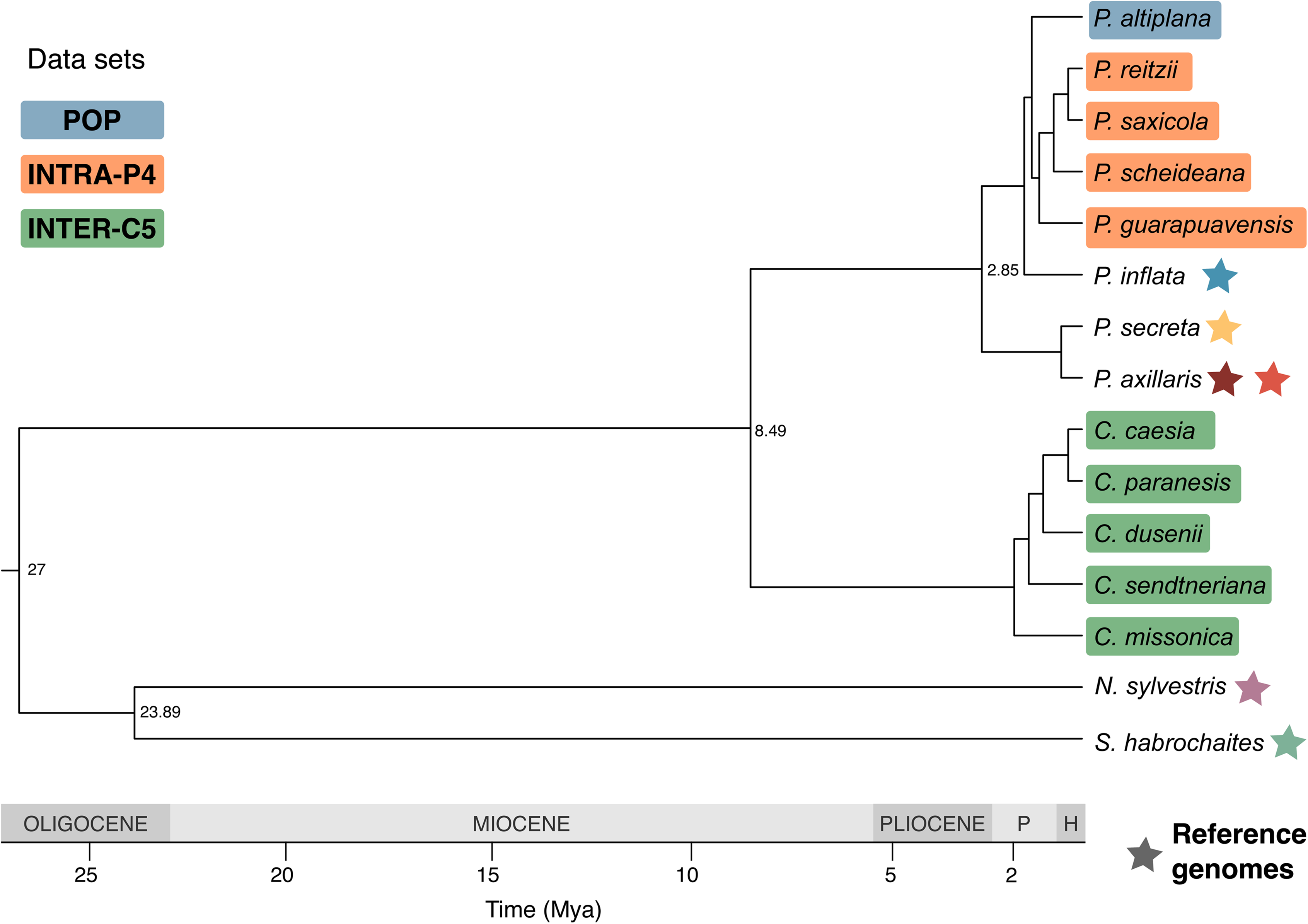

Each dataset was analyzed using six reference genomes that span a gradient of phylogenetic distances. Four belong to *Petunia* – *P. inflata* and *P. axillaris* I (Bombarely et al., 2016), *P. axillaris* II (https://www.ncbi.nlm.nih.gov/datasets/genome/GCA_029990575.1/), and *P. secreta* (https://www.ncbi.nlm.nih.gov/datasets/genome/GCA_029990175.1/). *Petunia inflata* represents the available closest reference for the POP and INTRA-P4 datasets, whereas *P. axillaris* and *P. secreta* belong to a different clade and serve as more distant congeneric references (Figure 1). We chose to use two *P. axillaris* genomes as they represent different sequencing and assembly methodologies. Specifically, *P. axillaris* I is a highly fragmented draft assembly generated solely from short reads from Illumina, while *P. axillaris* II results from long-read assemblies at the pseudo-molecule level, thus allowing us to assess how assembly quality affects SNP calling.

To evaluate how deeper phylogenetic divergence affects reference genome choice, we also included two additional Solanaceae genomes, *Nicotiana sylvestris* (Sierro et al., 2024) and *Solanum habrochaites* (Yu et al., 2022). These lineages last shared a common ancestor with the *Petunia*-*Calibrachoa* clade approximately 63.5 Mya (Huang et al., 2023). For comparison, we also used a *de novo* approach with STACKS v2.62 (Rivera-Colón & Catchen, 2022).

### SNP Calling and Filtering

The workflow for the analyses is illustrated in Figure 2. For the reference-based analyses, all sequence groups were processed identically for SNP calling. Raw reads were inspected with FASTQC v0.11.7 (Andrews, 2010) and MULTIQC (Ewels et al., 2016). Adapters and barcodes were removed, low-quality regions (< Q30) were trimmed, and reads < 50 bp were discarded with FASTQ-MCF v1.04.807 (Aronesty, 2013). Filtered reads were mapped to all reference genomes using the default settings of BWA v0.7.10-r789 (Li & Durbin, 2010). To assess whether the proportion of mapped reads differed among reference genomes, we used a linear mixed-effects model (mapping rate ∼ reference genome | samples). For post hoc pairwise comparisons, we estimated marginal means (EMMs) using Tukey’s correction for multiple testing. We used the *lme4* (Bates et al., 2015), *lmerTest* (Kuznetsova et al., 2017), and *emmeans* (Lenth & Piaskowski, 2025) packages in R (v4.3.0).

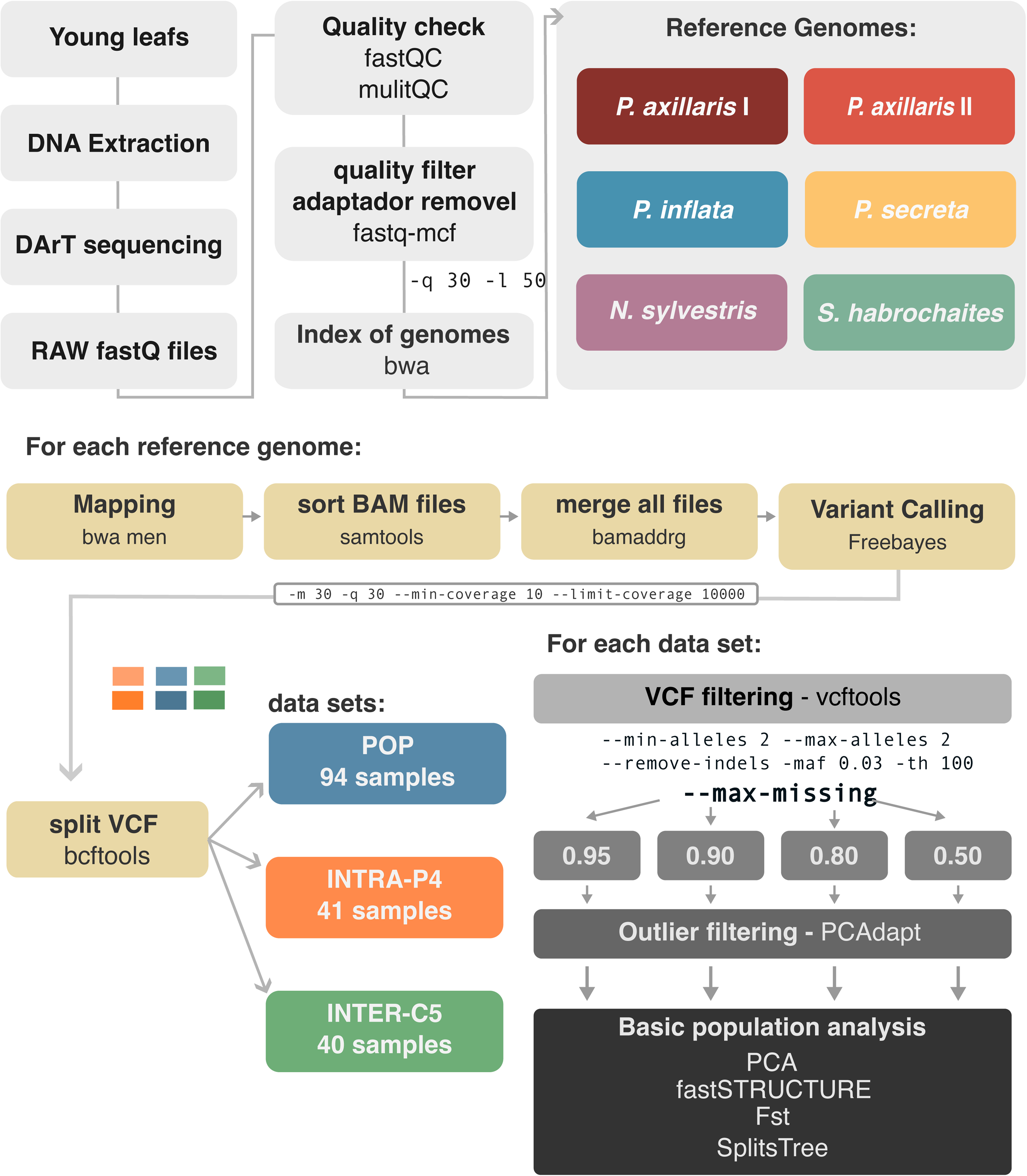

Unmapped reads were discarded, and BAM files were processed with SAMTOOLS v1.3.1 (Danecek et al., 2021). BAM files were merged per reference genome with bamaddrg (https://github.com/ekg/bamaddrg), sorted, and indexed with SAMTOOLS. Variants were called with FREEBAYES v1.3.6 (Garrison & Marth 2012) using mapping quality > 30, base quality > 30, and read depth > 10. Filtering thresholds varied by dataset. For the population-level group, SNPs with 10% missing data and minor allele frequency (--maf) ≥ 0.05 were included. Linkage disequilibrium (LD) was minimized by retaining only one SNP per 100 bp (--thin) using VCFTOOLS v0.1.12 (Danecek et al., 2011). We assessed the mapping rate (the proportion of total reads from each sample successfully mapped to the reference genome) and the number of SNPs recovered in each data group.

For the *de novo* approach, we cleaned the reads and removed barcodes using FASTQ-MCF, following the same procedure as in the reference-based pipeline. STACKS requires reads of the same length for *de novo* assembly, so all reads were trimmed to 100 bp. We used the *denovo_map.pl* module in STACKS to identify SNPs directly from the reads. Parameter optimization (Rivera-Colón & Catchen, 2022) was performed by running the *de novo* pipeline multiple times on a subset of 20 samples from each dataset, iterating over increasing values of M and n (from 1 to 9). This procedure aimed to determine the parameters that maximize the number of R80 loci—polymorphic loci present in at least 80% of samples. The optimal parameters (M = n = 3) were selected for the final *de novo* assembly and SNP calling. We filtered for missing data using the *population* STACKS module, retaining only loci present in at least 80% of individuals across populations (RL=L0.8), with a minor allele frequency (MAF) cutoff at 0.04, and we used only the first SNP of each read (—write-single-snp), preventing linkage disequilibrium.

### Genetic Diversity, Population Differentiation, and Phylogeny

We processed each data group separately and conducted standard population-genetics analyses to assess genetic diversity, structure, and phylogenetic relationships. We used the filtered datasets, which excluded monomorphic sites, to estimate genetic diversity indices, including nucleotide diversity (π, calculated per site), private alleles, polymorphic site proportion (P), observed (Ho) and expected (He) heterozygosity, inbreeding coefficients (*F*_IS_), and pairwise fixation index (*F*_ST_) using STACKS. Finally, we performed a principal components analysis (PCA) using the *gl.pcoa* function in DARTR (Mijangos et al., 2022).

Population and species structure were inferred using fastSTRUCTURE v1.074 (Raj et al., 2014). We ran K from 1 to 15 with 10 replicates per run. The STRUCTURE_THREADER software (Pina-Martins et al., 2017) was used, and the results were summarized using STRUCTURE HARVESTER (Earl & von Holdt, 2012). The most likely number of clusters was evaluated by inspecting likelihood plots and applying Evanno’s method (Evanno et al., 2005). We employed POPHELPER (Francis, 2017) to plot the graphs, organizing them primarily to maintain a consistent color pattern in the GRAPHIC software.

We used the NeighborNet method in SPLITSTREE v4.16 (Huson and Bryant, 2006) to construct a phylogenetic network for each dataset. We plotted and compared the network topologies to assess whether the choice of genome reference affects the primary evolutionary interpretation of the data.

### Does the Number of Reads Matter?

To evaluate the impact of sequencing depth on summary statistics, we simulated low-coverage sequencing by randomly subsampling reads from each sample. We used the POP and INTRA-P4 datasets for this analysis, along with custom Python and Bash scripts to generate subsampled datasets (https://github.com/luassoares/Compare-Reference-Genome). We tested 50,000, 100,000, 200,000, and up to 500,000 reads per sample to reflect realistic variation in genome coverage. Each subsampling scheme was replicated ten times to account for the stochastic variation introduced by the random sampling process. We mapped each subsampled dataset to the *P. axillaris* I genome. We evaluated the depth and breadth of coverage for each resampling using the SAMTOOLS *depth* command to obtain the depth coverage per site for all sites (including the unmapped) and then divided the number of sites with depth coverage > 10x by the total number of sites to get the proportion of covered genome (breadth coverage). Summary population statistics were calculated for each dataset, namely the number of private alleles, π, Ho, and *F*_IS_.

## Results

### Mapping Rates

The mapping rates were positively associated with phylogenetic proximity between the dataset and the reference genome (Figure 3). Across all datasets, mapping rates did not differ significantly among the *P. axillaris* I, *P. axillaris* II, and *P. secreta* reference genomes. For the congeneric datasets (POP and INTRA-P4), mapping to the closest genome, *P. inflata,* yielded slightly higher average mapping rates, which were significantly higher than those for the other congeneric references (Supplementary Tables S2 and S3, Figure 3).

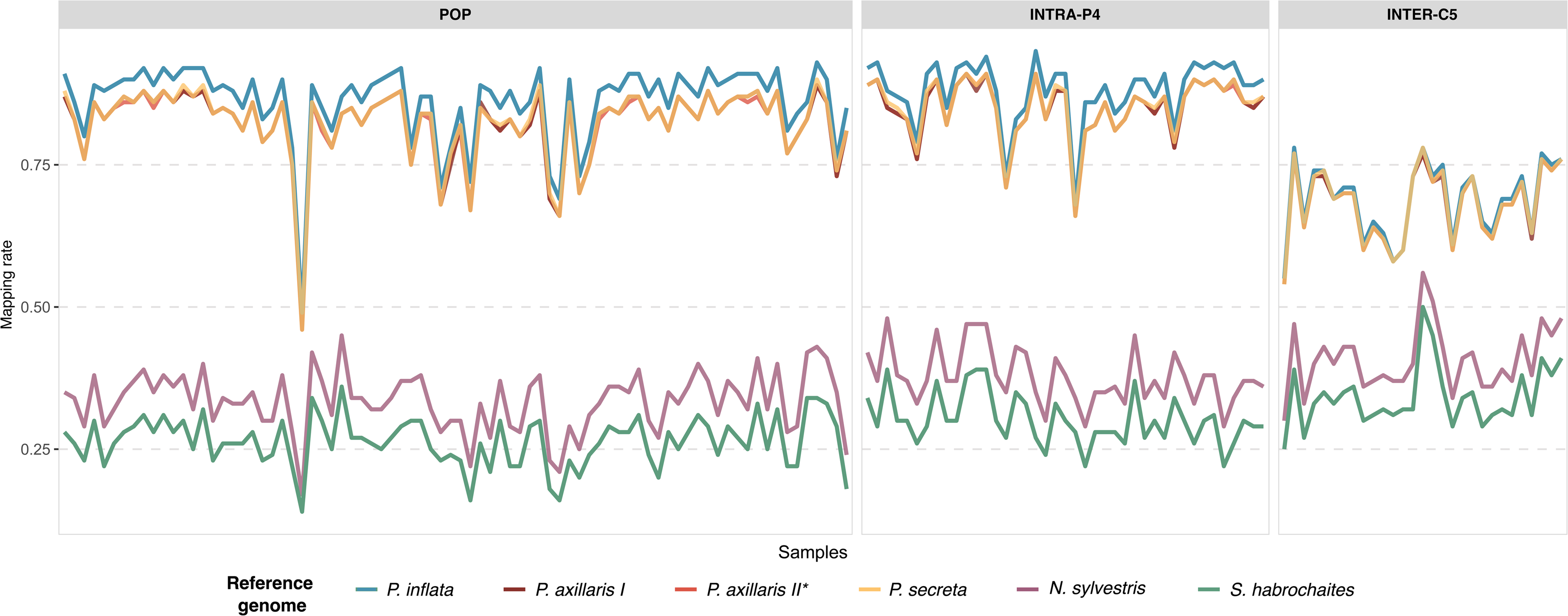

As expected, mapping rates decreased when using more distantly related reference genomes. *Nicotiana sylvestris* consistently showed higher mapping rates than *Solanum habrochaites*. The INTER-C5 dataset showed lower mapping rates than POP and INTRA-P4 when using *Petunia* genomes as a reference. In contrast, INTER-C5 showed slightly higher mapping rates when we used the most phylogenetically distant reference genomes, *Nicotiana* and *Solanum*.

### SNP Discovery

After mapping, SNP calling, and filtering, we generated 21 VCF files across three datasets, six reference genomes, and the *de novo* assembly. SNP recovery followed the same overall pattern across all datasets (Supplementary Figure S1). The four *Petunia* reference genomes consistently produced the highest SNP counts, which were nearly identical among them. The only exception was the POP dataset, for which *P. inflata* recovered slightly more SNPs than the other *Petunia* genomes. In contrast, mapping to the more distantly related references (*Nicotiana sylvestris* and *Solanum habrochaites*) produced markedly fewer SNPs, approximately 5- to 7-fold fewer variants. The *de novo* assembly yielded intermediate SNP counts, falling between those obtained with the congeneric *Petunia* references and those with distant genomes. This pattern was consistent across all datasets; however, for INTRA-P4, the *de novo* assembly performed poorly, retaining only 41 SNPs after filtering.

### Population level statistics

We analyzed summary statistics for all datasets across the reference genomes and the *de novo* approach (Supplementary Figure S2). The number of private alleles and Ho differed significantly among datasets, whereas π varied significantly only in the INTER-C5 dataset. Across all datasets, the number of private alleles was higher with closely related reference genomes than with distant ones, although no significant differences were observed among the *Petunia* genomes themselves. The more distant references yielded higher π values and, consequently, higher Ho. The *de novo* approach performed comparably to *Petunia* genomes in terms of private alleles in the INTER-C5 dataset, but its results were more similar to those obtained with distant references for the INTRA-P4 and POP datasets. Overall, it produced the lowest levels of π and Ho.

No statistically significant differences were detected in *F*_IS_, although the POP dataset tended to show slightly lower values. Although the statistical power was insufficient to detect significant differences in private alleles and Ho in the POP dataset, the graphical results clearly illustrate the distinct patterns among reference genome choices and analytical approaches (Supplementary Figure S2).

### Population Structure and Evolutionary Relationships

We assessed the population structure and evolutionary relationships for each VCF file. Despite differences in SNP counts across reference genomes, the overall results were highly congruent across all analyses, even for the most distant reference genomes.

PCA results were consistent across reference genomes (Figure 4). Although each principal component (PC) axis explained a slightly different percentage of variance, the relative positions of individuals and clusters remained stable throughout the analyses. Even for INTRA-P4, with only 41 SNPs, the PCA could recover the variation of the samples, even though it was not as good as the other. This finding indicates that population structure and evolutionary patterns were robust, regardless of the reference genome used.

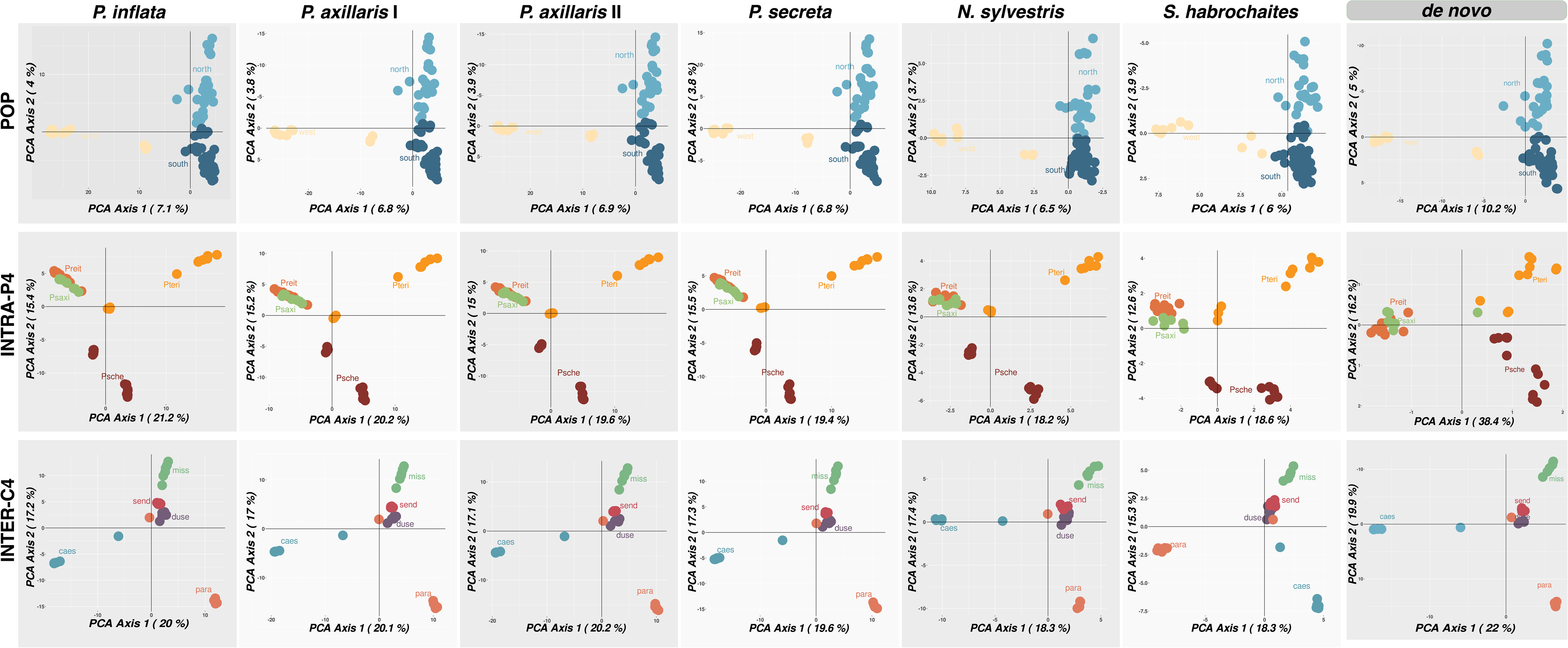

*F*_ST_ values were also consistent across reference genomes (Supplementary Figure S3), with no significant variation among them. The overall pattern of population differentiation and the relative relationships among groups remained stable. This result suggests that the biological interpretation is unaffected, whereas the reference genome might influence the magnitude of *F*_ST_.

fastSTRUCTURE results followed similar trends, although there was slightly more variation across reference genomes. As is typical with this method, no single best K was inferred. Instead, the analysis returned a range of plausible K values (Supplementary Table S4). INTRA-P4, mapped to the *P. axillaris* I reference genome, had the greatest variation in possible K (5 – 10). We selected the best K based on previous studies for each dataset to plot and explore.

Overall, the reference genomes and *de novo* approach produced consistent population structure patterns. For the POP dataset (Figure 5a), we identified the three geographical groups of *P. altiplana*. Most reference genomes yielded nearly identical results, except for the *P. secreta* genome, which suggested different ancestry proportions for several individuals. For INTRA-P4 (K = 5; Figure 5b), minor variation in ancestry components was observed when using the *P. axillaris* genome, reflecting the complex evolutionary relationships among populations rather than true reference bias. A similar pattern was observed for the INTER-C5 dataset (Figure 5c), where the best K = 5 corresponded to the number of species. In this case, population structure was largely consistent across references, with each species showing a distinct genetic component.

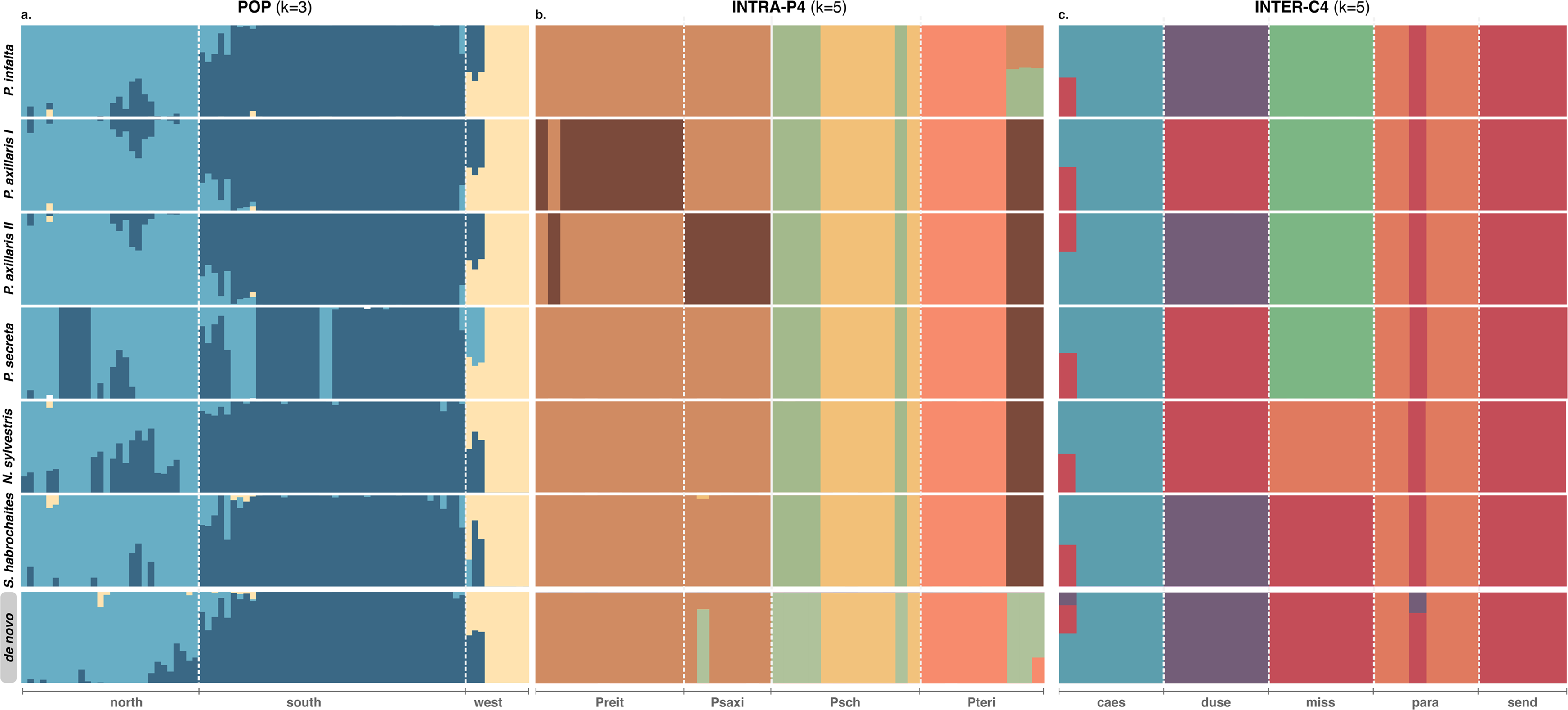

SPLITSTREE analyses (Figure 6) revealed consistent topologies across reference genomes. Trees based on the *Nicotiana* and *Solanum* genomes, as well as the *de novo* assembly, exhibited broader internal nodes, likely reflecting the reduced number of SNPs. Nevertheless, the overall relationships among individuals and groups remained consistent. Even for the INTRA-P4 *de novo* dataset, which retained only 41 SNPs, the phylogenetic relationships among species were correctly recovered, indicating that all approaches provided sufficient phylogenetic signal.

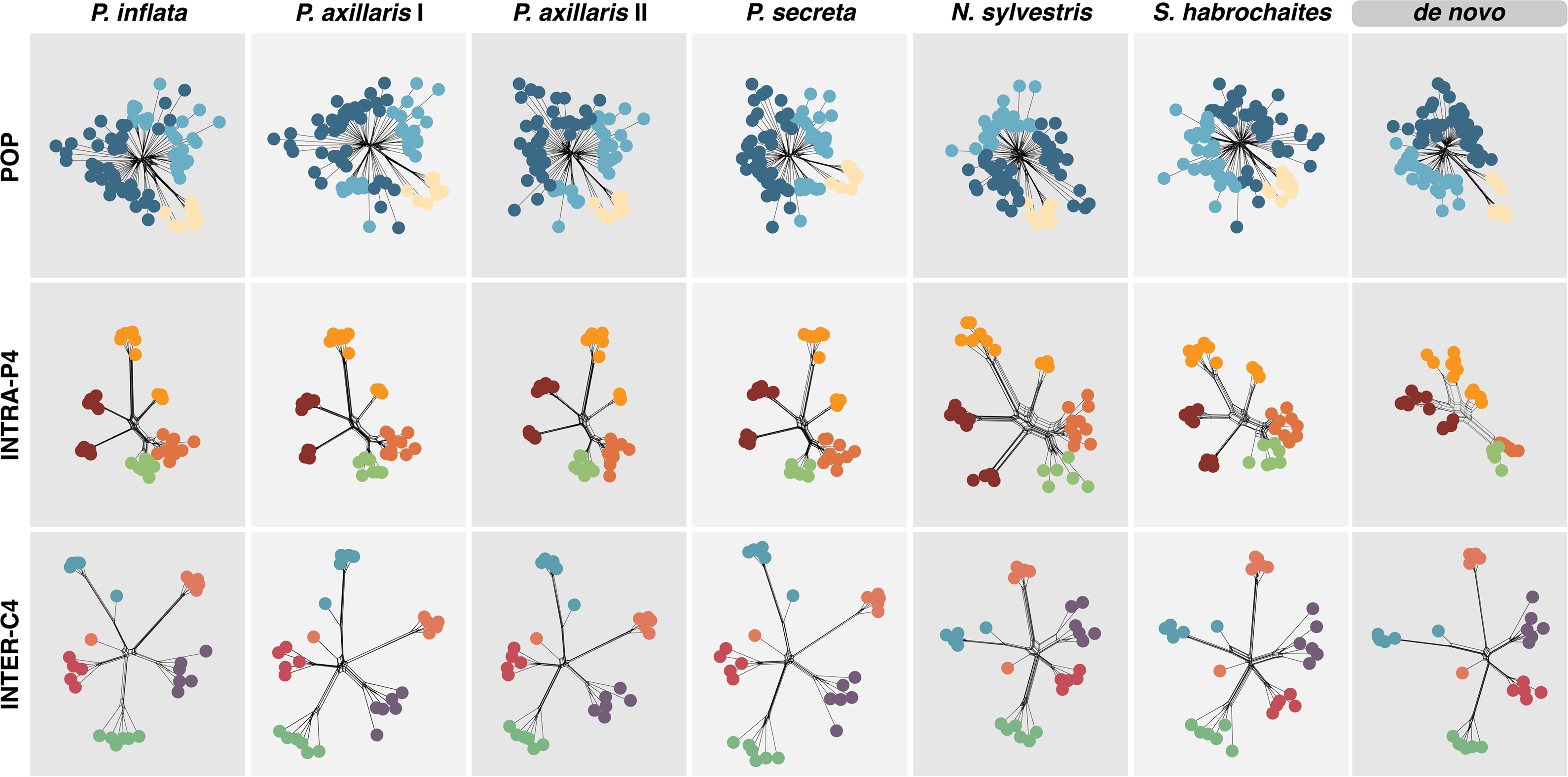

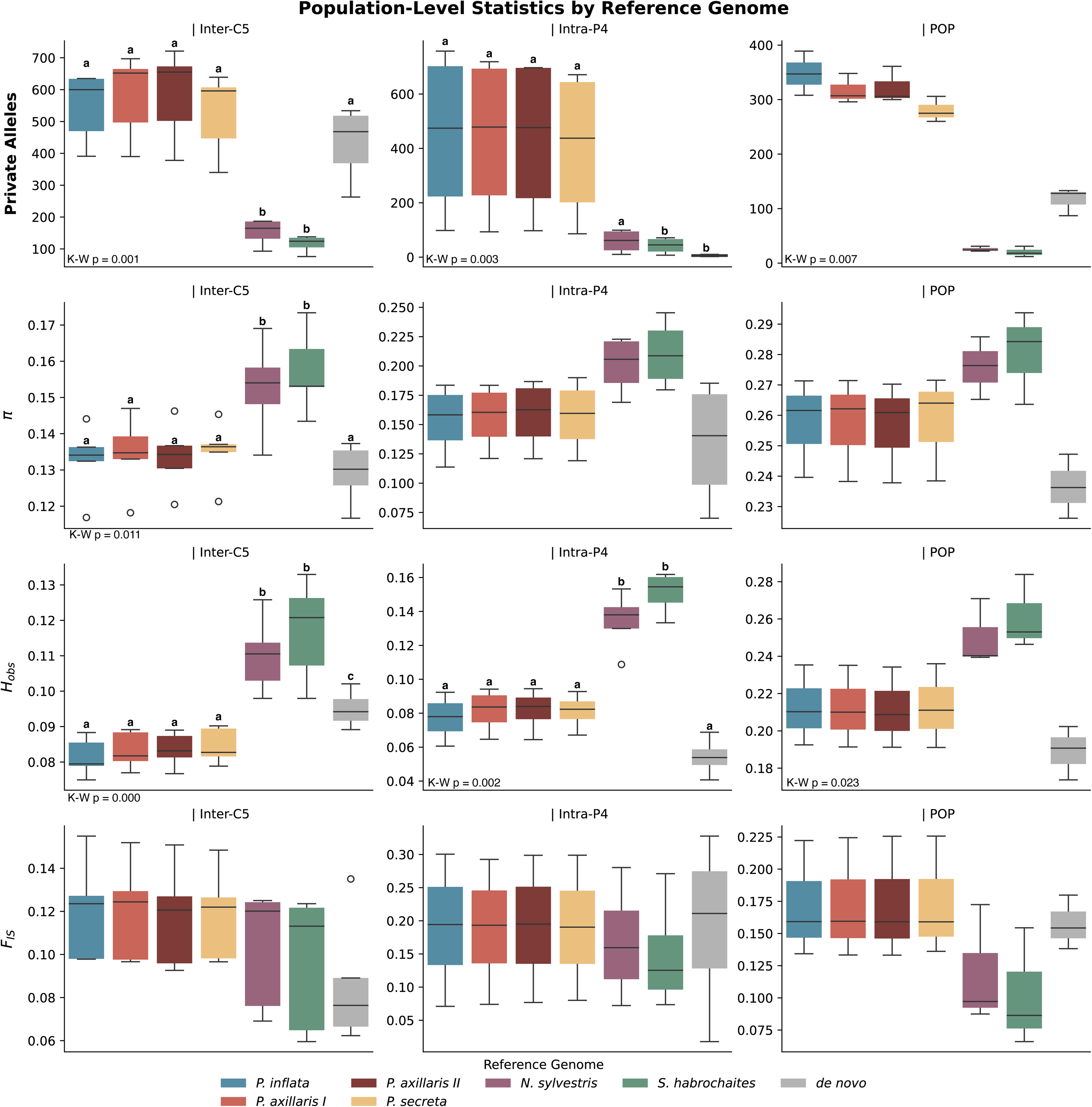

### Does the Number of Reads Matter?

Across the subsampled datasets, we observed variation in the absolute values of genetic diversity indices (Supplementary Figure S4). For the POP dataset, summary statistics showed higher estimates and greater variability among populations at different sequencing depths. In contrast, for INTRA-P4, the number of private alleles increased with increasing read counts and then stabilized at 200,000 reads, indicating that sequencing had reached a diversity plateau. Beyond 300,000 reads, most additional reads were shared across species, leading to a decline in Ho, π, and *F*_IS_ after 100,000 reads.

Despite these quantitative differences, the overall patterns and relative comparisons among populations remained consistent across subsampling schemes (Supplementary Figure S4). This suggests that, although sequencing depth influences the precision of diversity estimates, the general interpretation of population structure and diversity patterns is robust to moderate reductions in genome coverage.

## Discussion

Despite substantial methodological advances in generating and processing of genomic data, downstream population genomic inferences remain sensitive to several key factors, particularly sequencing depth and the choice of a reference genome for variant calling (Andrews et al., 2016). In this study, we evaluated how these factors influence estimates of genetic diversity, population structure, and evolutionary relationships in recently and rapidly diversified plant lineages. Using DArT-derived SNPs from multiple *Petunia* species and related taxa as a model system, we compared the effects of mapping reads to both closely and distantly related reference genomes. When using congeneric reference genomes, we observed highly consistent mapping rates, SNP recovery, and downstream population genomic patterns. In contrast, mapping to the more distantly related species resulted in lower mapping rates and stronger effects on summary statistics. However, despite these quantitative reductions, the broader results for genetic structure, diversity (PCA and STRUCTURE), and evolutionary relationships (SPLITSTREE topology) remained largely congruent across reference genomes.

### Number of reads

In population-scale RAD-seq datasets, the number of reads per sample often varies due to technical and biological factors, leading to differences in effective genome coverage. Such variation can strongly influence downstream population genomic inferences. Loci with insufficient depth or missing haplotypes can produce summary statistics that deviate from true values (Arnold et al., 2013), and the inherently higher error rates of NGS compared to Sanger sequencing necessitate minimum depth thresholds to ensure reliable genotype calls (Huang & Knowles, 2016). In our tests, we observed that as sequencing depth increases, the number of recoverable loci and the amount of phylogenetic signal also increase, but only up to a point (Supplementary Figure S4). For example, in the INTRA-P4 dataset, a congeneric dataset, private allele counts reached a plateau at ∼200,000 reads. After this point, additional reads contributed mainly to variation shared among the species, reflected in declining He (Supplementary Figure S4). This finding indicates that additional sequencing does not always increase the amount of informative variation for population analyses. In contrast, the POP dataset, which is a population dataset from a highly diverse species (Soares et al., 2023, 2024) did not reach a plateau. In this case, more reads continued to yield new polymorphisms. These observations indicate that the optimal read depth depends on the biological characteristics of the system, sample size, and experimental design. Even so, datasets with lower coverage can still provide insights into evolutionary relationships. A reduction in matrix size that results from stricter depth filters should not be interpreted as evidence that higher coverage is always necessary. Early benchmarks indicated that a minimum of three reads for homozygous loci and four reads for heterozygous loci are required to achieve 95% genotype accuracy, corresponding to a practical lower limit of about 5-fold coverage (Hohenlohe et al., 2010; Xu et al., 2019).

### Mapping

Mapping rates were strongly associated with the evolutionary distance of the reference genome: the closer the reference, the higher the proportion of aligned reads (Figure 3). This pattern is well documented, with closely related references consistently yielding higher mapping rates and more distant references producing lower rates (Günther & Nettelblad, 2019; Prasad et al., 2022). Alignments to distantly related genomes typically generate a greater proportion of low-scoring reads, whereas closely related references preferentially retain conserved regions with lower mutation rates after filtering (McCormack et al., 2009; Huang & Knowles, 2016). Moreover, DNA fragments carrying alleles identical to the reference sequence are more likely to map successfully and pass post-alignment quality thresholds, resulting in their overrepresentation in downstream analyses (Brandt et al., 2015; Ros-Freixedes et al., 2018).

When genomes have had more time to evolve independently, extensive structural rearrangements, documented in lineages that diverged during more than 20 million years (Moran et al., 2020; Reid et al., 2021), can exacerbate the challenges of using heterospecific reference genomes. In contrast, for recent radiations without large-scale genome duplications, the impact of reference choice is reduced. Consistent with this expectation, we observed stronger effects of reference genome divergence in distantly related genomes than in congeneric *Petunia* references. Although in the most recent phylogeny *Nicotiana* and *Solanum* are similarly distant from *Petunia* (Deanna et al., 2026), the *Nicotiana* genome consistently produced higher alignment rates than *Solanum* (Figure 3). Assembly of reads to a reference genome is based on sequence similarity (Catchen et al., 2013), and reads with higher mutation rates and higher variability will tend to have lower alignment scores (Nielsen et al., 2011; Huang & Knowles, 2016). So, this could reflect differences in genome structural evolution, with *Solanum* exhibiting a higher degree of genome rearrangement or divergence from *Petunia* than *Nicotiana* (Frary et al., 2016; Deanna et al., 2022).

The choice of reference genome had a marked effect on SNP recovery and summary statistics across all datasets (Supplementary Figures S1, S2). Mapping to different congeneric reference genomes yielded similar SNP recovery and population genetic estimates, whereas using more distantly related references resulted in reduced SNP counts and increased variability in summary statistics. This is consistent with previous studies documenting reference genome bias in population genomic analyses (Shafer et al., 2017; Günther & Nettelblad, 2019; Reid et al., 2021; Prasad et al., 2022). The *de novo* approach recovered more SNPs than the distant references but exhibited the greatest variability in summary statistics (Supplementary Figures S1, S2). Notably, these quantitative differences did not necessarily translate into major effects on downstream analyses, supporting earlier findings that a higher number of polymorphic loci does not always yield stronger or more stable signals of population differentiation (Díaz-Arce & Rodríguez-Ezpeleta, 2019). However, these changes may affect any analysis associated with quantitative genetics that relies on variant density (e.g., GWAS, LD; van den Berg et al., 2019).

The performance of a reference genome for SNP discovery is closely tied to the evolutionary history and structural similarity between the reference and target genomes. In taxa with relatively conserved karyotypes, such as birds, heterospecific references can perform well in SNP discovery (Ellegren, 2010; Galla, 2019). In contrast, lineages characterized by rapid chromosomal evolution often show reduced mapping quality and increased bias when distant references are used (Vershinina & Lukhtanov, 2017). *Petunia* species diverged less than two million years ago and show no evidence of major structural genome changes, which likely explains the similar SNP recovery observed across all *Petunia* reference genomes. In line with these findings, our results show a much stronger reduction in SNP recovery when using distant reference genomes compared to congeneric references.

Despite quantitative differences in SNP counts and summary statistics, PCA, STRUCTURE, and phylogenetic analyses revealed broadly consistent patterns across reference genomes, with only minor variation among methods and datasets (Figures 4, 5, and 6). PCA is generally less sensitive to reference bias because it captures broad axes of genomic variation rather than relying on specific alignment characteristics (Rick et al., 2024). Similar patterns, characterized by substantial quantitative differences in mapping rates and SNP counts but consistent qualitative results in diversity and structure, have been reported in studies evaluating reference bias using different cultivars or strains (Ahmad et al., 2025). Likewise, comparisons between dog and wolf reference genomes have shown that reference choice can affect coverage, variant recovery, and heterozygosity estimates, while overall population structure remains stable (Gopalakrishnan et al., 2017).

Phylogenetic analyses can, in some cases, be more sensitive to reference choice, as small changes in alignment or SNP retention may influence topology or clade resolution (Rick et al., 2024). In our analyses, these effects were subtle and became more apparent only when the most distantly related reference genomes were used (Figure 6). Similar robustness has been reported in *Mycobacterium*, where phylogenetic trees remained stable across all but the most divergent reference genomes, indicating that reference choice can be flexible when lineage divergence is shallow (Lee & Behr, 2016). This pattern mirrors the evolutionary scale of the taxa examined here. Importantly, regardless of whether congeneric or more distant reference genomes were used, the overall patterns of population differentiation and the interpretation of phylogenetic relationships remained consistent.

The effects of reference genome choice may be more pronounced in analyses that target fine-scale genomic features, which were not explored in this study. Divergent reference genomes have been shown to bias inferences of demographic history, recombination landscapes, and ancient genome reconstructions (Slabaugh et al., 2019; Akopyan et al., 2025). In addition, missing data can affect downstream population genomic analyses, particularly when combined with reference bias. Missing data is a well-studied issue, and general recommendations favor minimizing missingness where possible (Hovmölleret al., 2013; Chattopadhyay et al., 2014; Marandel, 2020; Hemstrom et al., 2025). We did not explicitly evaluate its interaction with reference genome choice here.

Overall, our results indicated that reference choice matters most when genomes are distantly related or, more likely, when analyses target fine-scale genomic signals. For recent radiations such as *Petunia* and *Calibrachoa*, where genome structure is largely conserved, closely related reference genomes yield comparable SNP datasets and minimize downstream bias. Using a congeneric genome as a reference yields the same biological conclusions about population structure and phylogenetic relationships.

These findings have important implications for studies of non-model species. For closely related taxa that diverged recently, a congeneric reference genome is sufficient for recovering population structure, phylogenetic relationships, and downstream population genomic patterns. When no closely related reference genome is available, more distantly related genomes can still provide useful information and may outperform *de novo* approaches, provided that a sufficient number of informative loci are recovered. Under these conditions, key evolutionary relationships can be inferred reliably, contributing to a deeper understanding of non-model species even in the absence of optimal genomic resources.

## Data Accessibility

Raw sequence reads are deposited in the GenBank with accessions in the Supplementary material. The VCF files are available at https://doi.org/10.6084/m9.figshare.31157242

## Benefit-Sharing

Benefits Generated: Benefits from this research accrue from the sharing of our data and results on public databases as described above.

## Author Contributions

LBF and LSS designed the research, LSS, LTG, and SG-R analyzed data, LBF and AB contributed analytical tools, LSS and LBF wrote the paper, and all authors read and approved the manuscript.

