## Supplementary Information for "Reference genome choice impacts SNP recovery but not evolutionary inference in young species"

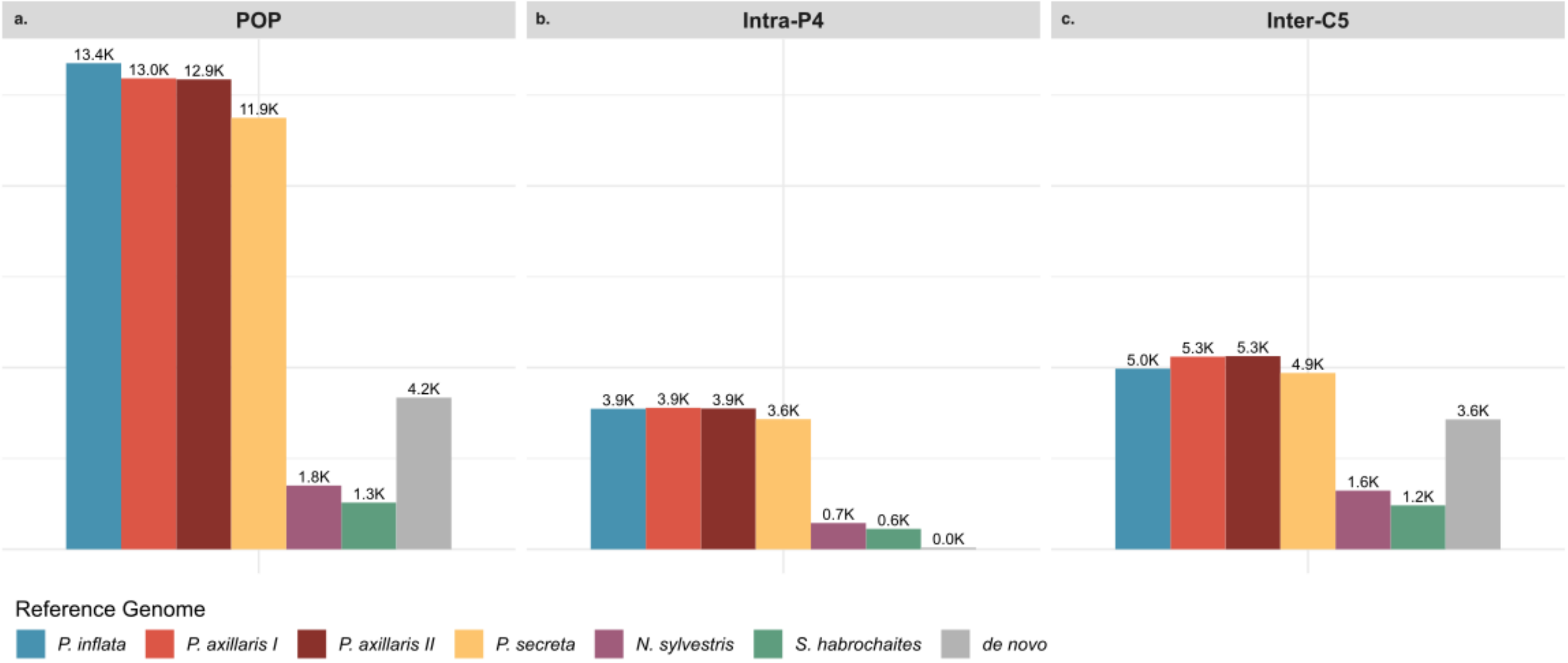


**Supplementary Figure S1.** Number of SNPs recovered across all datasets, reference genomes, and missing data thresholds applied during filtering.

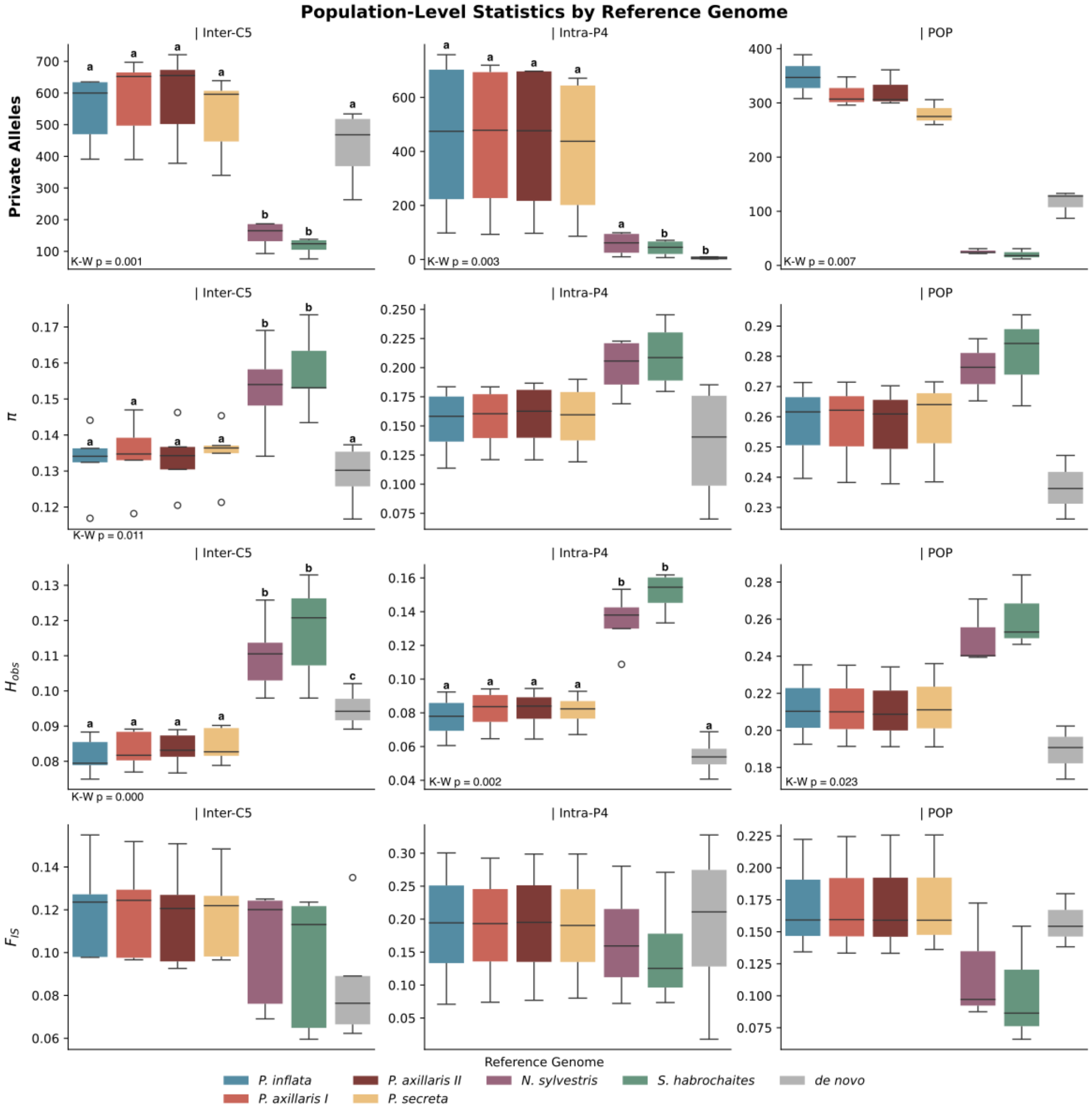

Population-Level Statistics by Reference Genome
| Inter-C5
| Intra-P4
| POP
Private Alleles
π
$H_{obs}$
$F_{IS}$
K-W p = 0.001
K-W p = 0.003
K-W p = 0.007
K-W p = 0.011
K-W p = 0.000
K-W p = 0.002
K-W p = 0.023
Reference Genome
P. inflata
P. axillaris I
P. axillaris II
P. secreta
N. sylvestris
S. habrochaites
de novo

**Supplementary Figure S2.** Population-level genetic statistics [number of private alleles, nucleotide diversity (π), observed heterozygosity (Ho), and inbreeding coefficient ($F_{IS}$)] across reference genomes. Reference genome abbreviations: Infl — *P. inflata*, AxiI — *P. axillaris* I, AxiII — *P. axillaris* II, Secr – *P. secreta*, Nico — *Nicotiana sylvestris*, Habro — *Solanum habrochaites*.

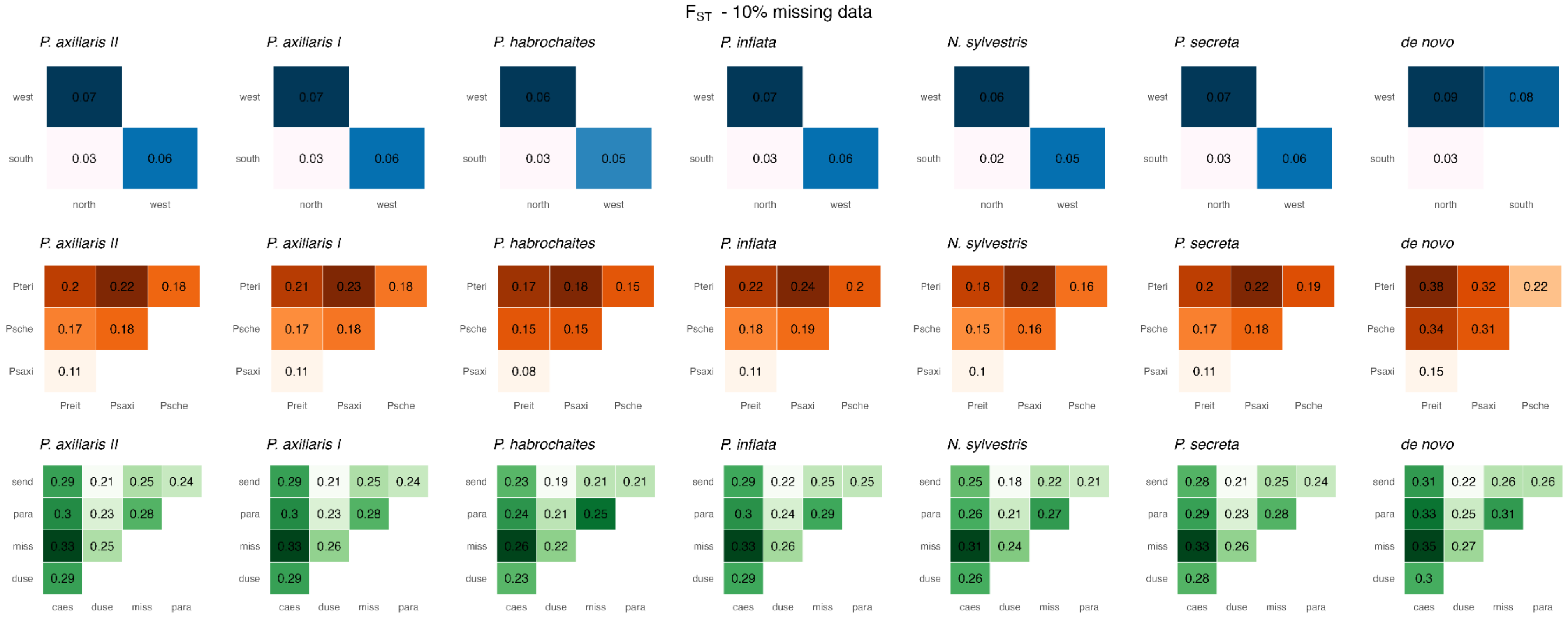


**Supplementary Figure S3** – Pairwise $F_{ST}$ estimates by dataset and reference genome, summarizing genetic differentiation across population and species groups under each alignment scenario. Shades of blue: POP—population-level dataset (80 *P. altiplana* individuals); Shades of orange: INTRA-P4—intraspecific dataset (41 individuals of four *Petunia* species); Shades of green: INTER-C5—interspecific dataset (40 individuals of five *Calibrachoa* species).

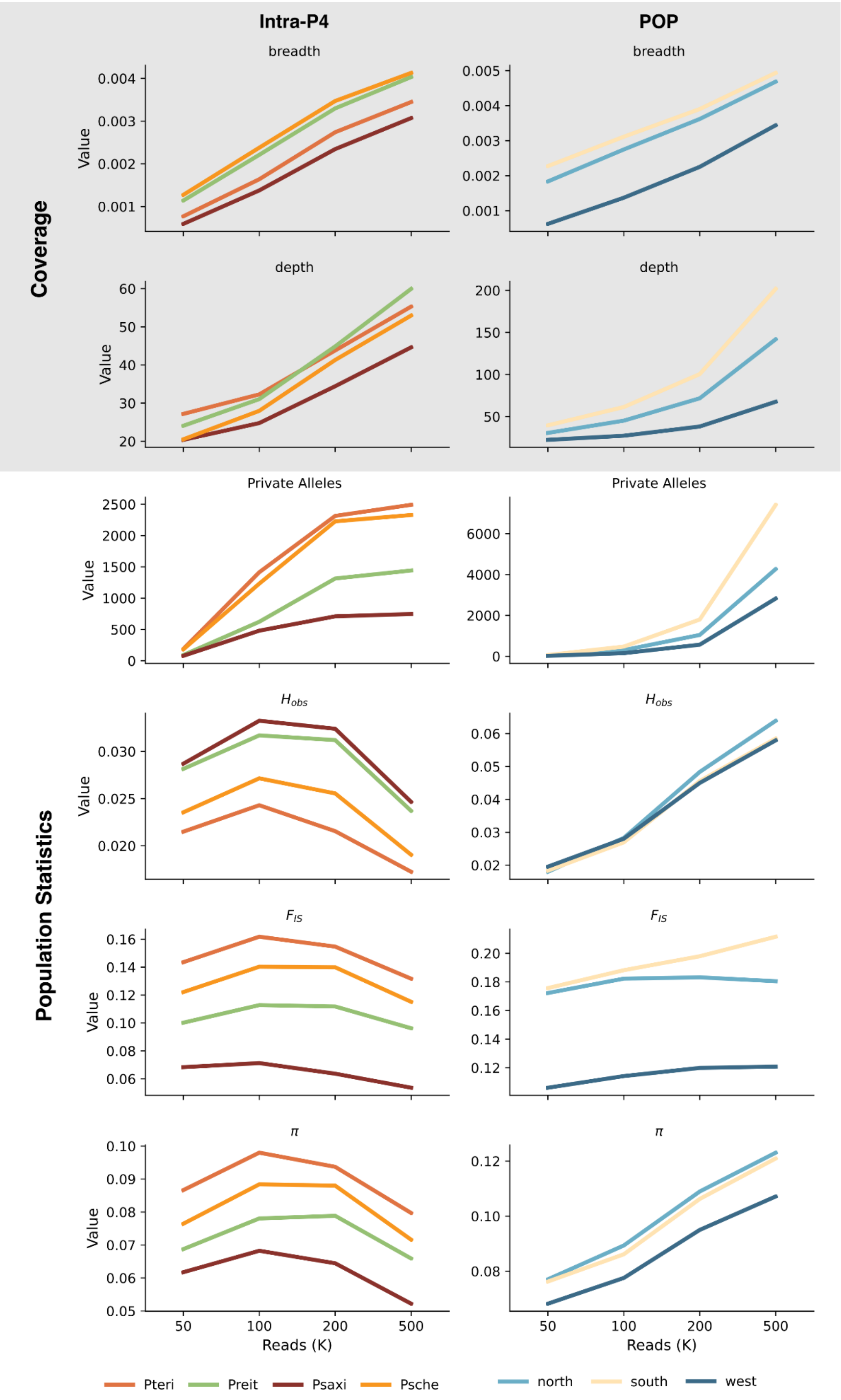


**Supplementary Figure S4.** Summary statistics, including the number of private alleles, observed heterozygosity (Ho), inbreeding coefficient ($F_{\text{IS}}$), and nucleotide diversity (π) for each dataset under different levels of subsampled read counts. Bars are colored by dataset.

**Supplementary Table S1.** GenBank accession and reads information.

| GenBank accession | Species | Group | Population | Sample | Raw reads | Sequence length raw (pb) | Filtered reads | Sequence length filtered (bp) |
|---|---|---|---|---|---|---|---|---|
| SRR29889316 | *Petunia reitzii* | Intra-P4 | Preit | 3005842 | 644150 | 138 | 307275 | 60 |
| SRR29889315 | *Petunia reitzii* | Intra-P4 | Preit | 3005924 | 439137 | 138 | 256320 | 60 |
| SRR29889314 | *Petunia reitzii* | Intra-P4 | Preit | 3006949 | 317025 | 138 | 194514 | 60 |
| SRR29889324 | *Petunia reitzii* | Intra-P4 | Preit | 3005899 | 595672 | 138 | 369795 | 60 |
| SRR29889313 | *Petunia reitzii* | Intra-P4 | Preit | 3005912 | 566754 | 138 | 409079 | 60 |
| SRR29889312 | *Petunia reitzii* | Intra-P4 | Preit | 3005852 | 616687 | 138 | 379782 | 60 |
| SRR29889322 | *Petunia reitzii* | Intra-P4 | Preit | 3006912 | 453621 | 138 | 263796 | 60 |
| SRR29889311 | *Petunia reitzii* | Intra-P4 | Preit | 3005888 | 551005 | 138 | 345935 | 60 |
| SRR29889323 | *Petunia reitzii* | Intra-P4 | Preit | 3005900 | 686099 | 138 | 390555 | 60 |
| SRR29889302 | *Petunia reitzii* | Intra-P4 | Preit | 3005911 | 564996 | 138 | 363170 | 60 |
| SRR29889291 | *Petunia reitzii* | Intra-P4 | Preit | 3005891 | 550804 | 138 | 328631 | 60 |
| SRR29889317 | *Petunia reitzii* | Intra-P4 | Preit | 3006944 | 557222 | 138 | 277223 | 60 |
| SRR29889308 | *Petunia saxicola* | Intra-P4 | Psaxi | 3005889 | 596950 | 138 | 354448 | 60 |
| SRR29889307 | *Petunia saxicola* | Intra-P4 | Psaxi | 3005901 | 568754 | 138 | 282926 | 60 |
| SRR29889306 | *Petunia saxicola* | Intra-P4 | Psaxi | 3005903 | 542861 | 138 | 286403 | 60 |
| SRR29889310 | *Petunia saxicola* | Intra-P4 | Psaxi | 3005877 | 627441 | 138 | 254155 | 60 |
| SRR29889305 | *Petunia saxicola* | Intra-P4 | Psaxi | 3005841 | 663313 | 138 | 406623 | 60 |
| SRR29889304 | *Petunia saxicola* | Intra-P4 | Psaxi | 3005853 | 624327 | 138 | 376725 | 60 |
| SRR29889309 | *Petunia saxicola* | Intra-P4 | Psaxi | 3005865 | 578902 | 138 | 281420 | 60 |
| SAMN40212237 | *Petunia scheideana* | Intra-P4 | Psche | 3005874 | 636778 | 138 | 370410 | 60 |
| SRR29889303 | *Petunia scheideana* | Intra-P4 | Psche | 3006945 | 294554 | 138 | 196766 | 60 |

| | | | | | | | | |
|---|---|---|---|---|---|---|---|---|
| SRR29889301 | *Petunia scheideana* | Intra-P4 | Psche | 2990934 | 405510 | 138 | 209465 | 60 |
| SRR29889300 | *Petunia scheideana* | Intra-P4 | Psche | 3005875 | 454332 | 138 | 300582 | 60 |
| SRR29889299 | *Petunia scheideana* | Intra-P4 | Psche | 3005887 | 544567 | 138 | 332408 | 60 |
| SRR29889297 | *Petunia scheideana* | Intra-P4 | Psche | 2990940 | 445755 | 138 | 148057 | 60 |
| SRR29889296 | *Petunia scheideana* | Intra-P4 | Psche | 3005851 | 490297 | 138 | 247600 | 60 |
| SAMN40212244 | *Petunia scheideana* | Intra-P4 | Psche | 3005886 | 499049 | 138 | 292913 | 60 |
| SAMN40212246 | *Petunia scheideana* | Intra-P4 | Psche | 3005839 | 571392 | 138 | 376961 | 60 |
| SRR29889298 | *Petunia scheideana* | Intra-P4 | Psche | 3006940 | 433443 | 138 | 251605 | 60 |
| SAMN40212236 | *Petunia scheideana* | Intra-P4 | Psche | 3005863 | 619624 | 138 | 378801 | 60 |
| SAMN54916668 | *Petunia scheideana* | Intra-P4 | Psche | 3006939 | 560323 | 138 | 345990 | 60 |
| SRR29889292 | *Petunia interior* | Intra-P4 | Pteri | 3005890 | 587270 | 138 | 333262 | 60 |
| SRR29889289 | *Petunia interior* | Intra-P4 | Pteri | 3005914 | 594035 | 138 | 307826 | 60 |
| SRR29889290 | *Petunia interior* | Intra-P4 | Pteri | 3005902 | 590091 | 138 | 287635 | 60 |
| SRR29889293 | *Petunia interior* | Intra-P4 | Pteri | 3005926 | 560907 | 138 | 289159 | 60 |
| SRR29889288 | *Petunia interior* | Intra-P4 | Pteri | 3005843 | 597476 | 138 | 283413 | 60 |
| SRR29889294 | *Petunia interior* | Intra-P4 | Pteri | 3005855 | 534488 | 138 | 317876 | 60 |
| SRR29889321 | *Petunia interior* | Intra-P4 | Pteri | 3005867 | 601252 | 138 | 312516 | 60 |
| SRR29889318 | *Petunia interior* | Intra-P4 | Pteri | 3005854 | 622692 | 138 | 356766 | 60 |
| SRR29889319 | *Petunia interior* | Intra-P4 | Pteri | 3006928 | 602569 | 138 | 362195 | 60 |
| SRR29889320 | *Petunia interior* | Intra-P4 | Pteri | 3006934 | 421631 | 138 | 270483 | 60 |
| SRR23348539 | *Petunia altiplana* | POP | north | 2062343 | 1627460 | 84 | 1043505 | 60 |
| SRR23348548 | *Petunia altiplana* | POP | north | 2062150 | 1892062 | 84 | 1188173 | 60 |
| SRR23348547 | *Petunia altiplana* | POP | north | 2062151 | 1949009 | 84 | 1285466 | 60 |
| SRR23348546 | *Petunia altiplana* | POP | north | 2062152 | 1566527 | 84 | 1002874 | 60 |
| SRR23348543 | *Petunia altiplana* | POP | north | 2062154 | 1744975 | 84 | 1140308 | 60 |
| SRR23348590 | *Petunia altiplana* | POP | north | 2062164 | 1494776 | 84 | 969290 | 60 |
| SRR23348589 | *Petunia altiplana* | POP | north | 2062165 | 1608830 | 84 | 1005467 | 60 |
| SRR23348555 | *Petunia altiplana* | POP | north | 2062166 | 2151254 | 84 | 1318466 | 60 |
| SRR23348544 | *Petunia altiplana* | POP | north | 2062167 | 2173840 | 84 | 1330427 | 60 |

| | | | | | | | | |
|---|---|---|---|---|---|---|---|---|
| SRR23348511 | *Petunia altiplana* | POP | north | 2062168 | 2173612 | 84 | 1336253 | 60 |
| SRR23348500 | *Petunia altiplana* | POP | north | 2062169 | 1761986 | 84 | 1083145 | 60 |
| SRR23348489 | *Petunia altiplana* | POP | north | 2062170 | 1549700 | 84 | 987680 | 60 |
| SRR23348577 | *Petunia altiplana* | POP | north | 2062171 | 1707138 | 84 | 1108084 | 60 |
| SRR23348588 | *Petunia altiplana* | POP | north | 2062172 | 1613844 | 84 | 999773 | 60 |
| SRR23348564 | *Petunia altiplana* | POP | north | 2062173 | 1606363 | 84 | 979280 | 60 |
| SRR23348563 | *Petunia altiplana* | POP | north | 2062174 | 2200728 | 84 | 1384700 | 60 |
| SRR23348562 | *Petunia altiplana* | POP | north | 2062175 | 2229221 | 84 | 1429395 | 60 |
| SRR23348561 | *Petunia altiplana* | POP | north | 2062176 | 1925787 | 84 | 1238328 | 60 |
| SRR23348560 | *Petunia altiplana* | POP | north | 2062177 | 1918502 | 84 | 1223622 | 60 |
| SRR23348559 | *Petunia altiplana* | POP | north | 2062178 | 1521224 | 84 | 1019558 | 60 |
| SRR23348558 | *Petunia altiplana* | POP | north | 2062179 | 1493440 | 84 | 984513 | 60 |
| SRR23348557 | *Petunia altiplana* | POP | north | 2062180 | 1550060 | 84 | 977017 | 60 |
| SRR23348556 | *Petunia altiplana* | POP | north | 2062181 | 1646145 | 84 | 986452 | 60 |
| SRR23348554 | *Petunia altiplana* | POP | north | 2062182 | 2070226 | 84 | 1311546 | 60 |
| SRR23348553 | *Petunia altiplana* | POP | north | 2062183 | 2165801 | 84 | 1370017 | 60 |
| SRR23348551 | *Petunia altiplana* | POP | north | 2062185 | 1717152 | 84 | 1114758 | 60 |
| SRR23348550 | *Petunia altiplana* | POP | north | 2062186 | 1435756 | 84 | 984664 | 60 |
| SRR23348549 | *Petunia altiplana* | POP | north | 2062187 | 1727978 | 84 | 1035076 | 60 |
| SRR23348537 | *Petunia altiplana* | POP | south | 2062188 | 1511202 | 84 | 981284 | 60 |
| SRR23348536 | *Petunia altiplana* | POP | south | 2062189 | 1444958 | 84 | 884754 | 60 |
| SRR23348534 | *Petunia altiplana* | POP | south | 2062191 | 2086484 | 84 | 1279922 | 60 |
| SRR23348532 | *Petunia altiplana* | POP | south | 2062192 | 1880107 | 84 | 1204591 | 60 |
| SRR23348527 | *Petunia altiplana* | POP | south | 2062196 | 1585698 | 84 | 1029422 | 60 |
| SRR23348524 | *Petunia altiplana* | POP | south | 2062198 | 1686569 | 84 | 1128217 | 60 |
| SRR23348523 | *Petunia altiplana* | POP | south | 2062199 | 1978938 | 84 | 1327767 | 60 |
| SRR23348521 | *Petunia altiplana* | POP | south | 2062200 | 2350379 | 84 | 1425058 | 60 |
| SRR23348519 | *Petunia altiplana* | POP | south | 2062202 | 1684194 | 84 | 1075914 | 60 |
| SRR23348516 | *Petunia altiplana* | POP | south | 2062204 | 982154 | 84 | 629148 | 60 |

| | | | | | | | | |
|---|---|---|---|---|---|---|---|---|
| SRR23348515 | *Petunia altiplana* | POP | south | 2062205 | 1643100 | 84 | 919608 | 60 |
| SRR23348514 | *Petunia altiplana* | POP | south | 2062206 | 2088945 | 84 | 1310644 | 60 |
| SRR23348513 | *Petunia altiplana* | POP | south | 2062207 | 2454436 | 84 | 1506151 | 60 |
| SRR23348510 | *Petunia altiplana* | POP | south | 2062208 | 2023906 | 84 | 1311987 | 60 |
| SRR23348509 | *Petunia altiplana* | POP | south | 2062209 | 1900983 | 84 | 1200171 | 60 |
| SRR23348508 | *Petunia altiplana* | POP | south | 2062210 | 1444642 | 84 | 963247 | 60 |
| SRR23348507 | *Petunia altiplana* | POP | south | 2062211 | 2001022 | 84 | 1187097 | 60 |
| SRR23348506 | *Petunia altiplana* | POP | south | 2062212 | 1866653 | 84 | 1068211 | 60 |
| SRR23348505 | *Petunia altiplana* | POP | south | 2062213 | 1428287 | 84 | 847494 | 60 |
| SRR23348504 | *Petunia altiplana* | POP | south | 2062215 | 1519886 | 84 | 878805 | 60 |
| SRR23348503 | *Petunia altiplana* | POP | south | 2062216 | 1727865 | 84 | 1194049 | 60 |
| SRR23348502 | *Petunia altiplana* | POP | south | 2062217 | 2227037 | 84 | 1371609 | 60 |
| SRR23348501 | *Petunia altiplana* | POP | south | 2062218 | 2150076 | 84 | 1286188 | 60 |
| SRR23348499 | *Petunia altiplana* | POP | south | 2062219 | 2060371 | 84 | 1104041 | 60 |
| SRR23348498 | *Petunia altiplana* | POP | south | 2062220 | 1738027 | 84 | 1076708 | 60 |
| SRR23348497 | *Petunia altiplana* | POP | south | 2062221 | 1949919 | 84 | 995891 | 60 |
| SRR23348496 | *Petunia altiplana* | POP | south | 2062222 | 2375042 | 84 | 1531561 | 60 |
| SRR23348495 | *Petunia altiplana* | POP | south | 2062223 | 2320800 | 84 | 1489056 | 60 |
| SRR23348494 | *Petunia altiplana* | POP | south | 2062224 | 2319531 | 84 | 1524511 | 60 |
| SRR23348493 | *Petunia altiplana* | POP | south | 2062225 | 2045918 | 84 | 1324290 | 60 |
| SRR23348492 | *Petunia altiplana* | POP | south | 2062226 | 2092494 | 84 | 1275406 | 60 |
| SRR23348487 | *Petunia altiplana* | POP | south | 2062230 | 1752960 | 84 | 1128953 | 60 |
| SRR23348585 | *Petunia altiplana* | POP | south | 2062231 | 2685495 | 84 | 1657353 | 60 |
| SRR23348584 | *Petunia altiplana* | POP | south | 2062232 | 2196651 | 84 | 1345697 | 60 |
| SRR23348583 | *Petunia altiplana* | POP | south | 2062233 | 2249408 | 84 | 1475435 | 60 |
| SRR23348582 | *Petunia altiplana* | POP | south | 2062234 | 1954566 | 84 | 1225833 | 60 |
| SRR23348581 | *Petunia altiplana* | POP | south | 2062235 | 2120365 | 84 | 1317948 | 60 |
| SRR23348580 | *Petunia altiplana* | POP | south | 2062236 | 1922573 | 84 | 1186124 | 60 |
| SRR23348579 | *Petunia altiplana* | POP | south | 2062237 | 1648852 | 84 | 1018728 | 60 |

| | | | | | | | | |
|---|---|---|---|---|---|---|---|---|
| SRR23348578 | *Petunia altiplana* | POP | south | 2062238 | 3342929 | 84 | 2099213 | 60 |
| SRR23348576 | *Petunia altiplana* | POP | south | 2062239 | 2894168 | 84 | 1682373 | 60 |
| SRR23348531 | *Petunia altiplana* | POP | south | 2070207 | 1448507 | 84 | 717951 | 60 |
| SRR23348575 | *Petunia altiplana* | POP | west | 2062240 | 1632825 | 84 | 984078 | 60 |
| SRR23348574 | *Petunia altiplana* | POP | west | 2062241 | 2340910 | 84 | 1438844 | 60 |
| SRR23348573 | *Petunia altiplana* | POP | west | 2062242 | 2043055 | 84 | 1228918 | 60 |
| SRR23348570 | *Petunia altiplana* | POP | west | 2062155 | 1660858 | 84 | 1068520 | 60 |
| SRR23348569 | *Petunia altiplana* | POP | west | 2062156 | 1578501 | 84 | 1018503 | 60 |
| SRR23348568 | *Petunia altiplana* | POP | west | 2062157 | 1570394 | 84 | 984926 | 60 |
| SRR23348567 | *Petunia altiplana* | POP | west | 2062158 | 2271090 | 84 | 1406341 | 60 |
| SRR23348491 | *Petunia altiplana* | POP | west | 2062227 | 1889036 | 84 | 1204787 | 60 |
| SRR23348490 | *Petunia altiplana* | POP | west | 2062228 | 1588468 | 84 | 997484 | 60 |
| SRR23348488 | *Petunia altiplana* | POP | west | 2062229 | 1759341 | 84 | 1094153 | 60 |
| SRR32113741, SRR32113742 | *Calibrachoa caesia* | Inter-C5 | caes | caes1 | 1107532 | 123 | 638960 | 60 |
| SRR32113676, SRR32113391 | *Calibrachoa caesia* | Inter-C5 | caes | caes12 | 885144 | 123 | 580032 | 60 |
| SRR32113422, SRR32113778 | *Calibrachoa caesia* | Inter-C5 | caes | caes3 | 1110527 | 123 | 565182 | 60 |
| SRR32113768, SRR32113518 | *Calibrachoa caesia* | Inter-C5 | caes | caes7 | 1031950 | 123 | 632731 | 60 |
| SRR32113529, SRR32113508 | *Calibrachoa caesia* | Inter-C5 | caes | caes8 | 1077151 | 123 | 670379 | 60 |
| SRR32113743, SRR32113729 | *Calibrachoa caesia* | Inter-C5 | caes | caes9 | 1110593 | 123 | 667921 | 60 |
| SRR32113390, SRR32113381 | *Calibrachoa dusenii* | Inter-C5 | duse | duse11 | 1031952 | 123 | 620715 | 60 |
| SRR32113555, SRR32113566 | *Calibrachoa dusenii* | Inter-C5 | duse | duse12 | 978047 | 123 | 600177 | 60 |
| SRR32113532, SRR32113543 | *Calibrachoa dusenii* | Inter-C5 | duse | duse2 | 1186463 | 123 | 508495 | 60 |

| | | | | | | | | |
|---|---|---|---|---|---|---|---|---|
| SRR32113665, SRR32113654, SRR32113643, SRR32113420 | *Calibrachoa dusenii* | Inter-C5 | duse | duse23 | 1747440 | 123 | 958123 | 60 |
| SRR32113421, SRR32113406 | *Calibrachoa dusenii* | Inter-C5 | duse | duse31 | 675979 | 123 | 365086 | 60 |
| SRR32113398, SRR32113782 | *Calibrachoa dusenii* | Inter-C5 | duse | duse7 | 1084977 | 123 | 614348 | 60 |
| SRR32113491, SRR32113490 | *Calibrachoa missionica* | Inter-C5 | miss | miss10 | 922675 | 123 | 504789 | 60 |
| SRR32113489, SRR32113488 | *Calibrachoa missionica* | Inter-C5 | miss | miss36 | 1014668 | 123 | 645919 | 60 |
| SRR32113602, SRR32113601 | *Calibrachoa missionica* | Inter-C5 | miss | miss38 | 1057728 | 123 | 566299 | 60 |
| SRR32113600, SRR32113599 | *Calibrachoa missionica* | Inter-C5 | miss | miss39 | 1101651 | 123 | 579809 | 60 |
| SRR32113598, SRR32113486 | *Calibrachoa missionica* | Inter-C5 | miss | miss40 | 957292 | 123 | 607259 | 60 |
| SRR32113485, SRR32113484 | *Calibrachoa missionica* | Inter-C5 | miss | miss58 | 975230 | 123 | 518262 | 60 |
| SRR32113589, SRR32113587 | *Calibrachoa paranensis* | Inter-C5 | para | para156 | 1061340 | 123 | 643798 | 60 |
| SRR32113586, SRR32113585 | *Calibrachoa paranensis* | Inter-C5 | para | para16 | 1082567 | 123 | 676620 | 60 |
| SRR36870411, SRR36870412 | *Calibrachoa paranensis* | Inter-C5 | para | para178 | 948955 | 123 | 531993 | 60 |
| SRR32113595, SRR32113594, SRR32113581, SRR32113582 | *Calibrachoa paranensis* | Inter-C5 | para | para227 | 1043273 | 123 | 617620 | 60 |
| SRR32113580, SRR32113579, SRR32113576, SRR32113578 | *Calibrachoa paranensis* | Inter-C5 | para | para240 | 1040292 | 123 | 631884 | 60 |

| | | | | | | | | |
|---|---|---|---|---|---|---|---|---|
| SRR32113575, SRR32113574 | *Calibrachoa paranensis* | Inter-C5 | para | para80 | 1069617 | 123 | 661153 | 60 |
| SRR32113551, SRR32113550 | *Calibrachoa sendtneriana* | Inter-C5 | send | send114 | 1051868 | 123 | 585169 | 60 |
| SRR32113549, SRR32113548, SRR32113546, SRR32113547 | *Calibrachoa sendtneriana* | Inter-C5 | send | send117 | 970330 | 123 | 565070 | 60 |
| SRR32113545, SRR32113544 | *Calibrachoa sendtneriana* | Inter-C5 | send | send121 | 980192 | 123 | 561134 | 60 |
| SRR32113559, SRR32113558, SRR32113557, SRR32113556 | *Calibrachoa sendtneriana* | Inter-C5 | send | send54 | 987972 | 123 | 558562 | 60 |
| SRR32113785, SRR32113567 | *Calibrachoa sendtneriana* | Inter-C5 | send | send9 | 900633 | 123 | 501661 | 60 |

**Supplementary Table S2.** Summary statistics for mapping rate using different reference genomes across all datasets.

| Grupo | Genome | variable | n | min | max | median | iqr | mean | sd | se | ci |
|---|---|---|---|---|---|---|---|---|---|---|---|
| POP | *N. sylvestris* | mapping | 80 | 0.17 | 0.45 | 0.34 | 0.07 | 0.334 | 0.052 | 0.006 | 0.012 |
| POP | *P. axillaris I* | mapping | 80 | 0.46 | 0.89 | 0.84 | 0.05 | 0.823 | 0.066 | 0.007 | 0.015 |
| POP | *P. axillaris II* | mapping | 80 | 0.46 | 0.89 | 0.84 | 0.05 | 0.824 | 0.067 | 0.007 | 0.015 |
| POP | *P. inflata* | mapping | 80 | 0.49 | 0.93 | 0.88 | 0.05 | 0.862 | 0.069 | 0.008 | 0.015 |
| POP | *P. secreta* | mapping | 80 | 0.46 | 0.9 | 0.84 | 0.05 | 0.825 | 0.067 | 0.007 | 0.015 |
| POP | *S. habrochaites* | mapping | 80 | 0.14 | 0.36 | 0.265 | 0.055 | 0.264 | 0.044 | 0.005 | 0.01 |
| INTRA-P4 | *N. sylvestris* | mapping | 41 | 0.29 | 0.48 | 0.37 | 0.06 | 0.378 | 0.049 | 0.008 | 0.015 |
| INTRA-P4 | *P. axillaris I* | mapping | 41 | 0.66 | 0.91 | 0.86 | 0.06 | 0.851 | 0.053 | 0.008 | 0.017 |
| INTRA-P4 | *P. axillaris II* | mapping | 41 | 0.66 | 0.91 | 0.86 | 0.06 | 0.851 | 0.053 | 0.008 | 0.017 |
| INTRA-P4 | *P. inflata* | mapping | 41 | 0.68 | 0.95 | 0.9 | 0.06 | 0.882 | 0.055 | 0.009 | 0.017 |
| INTRA-P4 | *P. secreta* | mapping | 41 | 0.66 | 0.91 | 0.86 | 0.06 | 0.853 | 0.052 | 0.008 | 0.017 |

| | | | | | | | | | | | |
|---|---|---|---|---|---|---|---|---|---|---|---|
| INTRA-P4 | *S. habrochaites* | mapping | 41 | 0.22 | 0.39 | 0.3 | 0.06 | 0.302 | 0.044 | 0.007 | 0.014 |
| INTER-C5 | *N. sylvestris* | mapping | 29 | 0.3 | 0.56 | 0.4 | 0.06 | 0.408 | 0.057 | 0.011 | 0.022 |
| INTER-C5 | *P. axillaris I* | mapping | 29 | 0.54 | 0.77 | 0.7 | 0.1 | 0.681 | 0.064 | 0.012 | 0.024 |
| INTER-C5 | *P. axillaris II* | mapping | 29 | 0.54 | 0.77 | 0.7 | 0.11 | 0.681 | 0.064 | 0.012 | 0.024 |
| INTER-C5 | *P. inflata* | mapping | 29 | 0.55 | 0.78 | 0.71 | 0.11 | 0.689 | 0.064 | 0.012 | 0.024 |
| INTER-C5 | *P. secreta* | mapping | 29 | 0.54 | 0.78 | 0.7 | 0.1 | 0.682 | 0.065 | 0.012 | 0.025 |
| INTER-C5 | *S. habrochaites* | mapping | 29 | 0.25 | 0.5 | 0.33 | 0.05 | 0.342 | 0.053 | 0.01 | 0.02 |

29

**Supplementary Table S3**. Estimated marginal means of mapping percentages across reference genomes for each dataset group.

| | Reference genome | emmean | SE | df | lower.CL | upper.CL | significance |
|---|---|---|---|---|---|---|---|
| **POP** | *S. habrochaites* | 0.264 | 0.007 | 108 | 0.250 | 0.278 | a |
| | *N. sylvestris* | 0.334 | 0.007 | 108 | 0.321 | 0.348 | b |
| | *P. axillaris I* | 0.823 | 0.007 | 108 | 0.809 | 0.836 | c |
| | *P. axillaris II* | 0.824 | 0.007 | 108 | 0.810 | 0.837 | c |
| | *P. secreta* | 0.825 | 0.007 | 108 | 0.811 | 0.839 | c |
| | *P. inflata* | 0.862 | 0.007 | 108 | 0.848 | 0.876 | d |
| **INTRA-P4** | *S. habrochaites* | 0.302 | 0.008 | 82.7 | 0.286 | 0.318 | a |
| | *N. sylvestris* | 0.378 | 0.008 | 82.7 | 0.362 | 0.394 | b |
| | *P. axillaris I* | 0.851 | 0.008 | 82.7 | 0.835 | 0.867 | c |
| | *P. axillaris II* | 0.851 | 0.008 | 82.7 | 0.835 | 0.867 | c |
| | *P. secreta* | 0.853 | 0.008 | 82.7 | 0.837 | 0.869 | c |
| | *P. inflata* | 0.882 | 0.008 | 82.7 | 0.866 | 0.898 | d |
| **INTER-C5** | *S. habrochaites* | 0.342 | 0.011 | 33.3 | 0.319 | 0.365 | a |
| | *N. sylvestris* | 0.408 | 0.011 | 33.3 | 0.385 | 0.431 | b |
| | *P. axillaris II* | 0.681 | 0.011 | 33.3 | 0.658 | 0.704 | c |
| | *P. axillaris I* | 0.681 | 0.011 | 33.3 | 0.658 | 0.705 | c |
| | *P. secreta* | 0.682 | 0.011 | 33.3 | 0.659 | 0.705 | c |
| | *P. inflata* | 0.689 | 0.011 | 33.3 | 0.666 | 0.712 | c |

Estimated marginal means (EMMeans) and associated standard errors (SE), degrees of freedom (df), and 95% confidence intervals (lower.CL and upper.CL) from linear mixed-effects models testing for differences in mapping percentages among reference genomes. Analyses were conducted separately for each dataset group (POP, INTRA-P4, and INTER-C5), with sample identity included as a random effect to account for repeated measures. Different letters in the "significance" column indicate statistically significant differences among reference genomes within each group (Tukey-adjusted $P < 0.05$).

**Supplementary Table S4.** K-values range estimated by fastSTRUCTURE analyses across reference genomes and datasets. Shown K are those that maximize marginal likelihood (model complexity), and the selected K-value to best describe the genetic structure in each dataset.

| **Reference genome** | bestK | | | | | |
|---|---|---|---|---|---|---|
| | **POP** | | **Intra-P4** | | **Inter-C5** | |
| *P. axillaris I* | 2 | 4 | 10 | 5 | 4 | 5 |
| *P. axillaris II* | 2 | 4 | 6 | 6 | 8 | 5 |
| *P. inflata* | 2 | 5 | 4 | 6 | 4 | 5 |
| *P. secreta* | 2 | 4 | 6 | 5 | 4 | 5 |
| *N. sylvestis* | 2 | 5 | 5 | 5 | 4 | 4 |
| *S. habrochaites* | 3 | 6 | 5 | 5 | 4 | 4 |
| *de novo* | 3 | 5 | 5 | 4 | 4 | 5 |

POP - population-level dataset (80 *P. altiplana* individuals); INTRA-P4 - intraspecific dataset (41 individuals of four *Petunia* species); INTER-C5 - interspecific dataset (40 individuals of five *Calibrachoa* species)
